# A deep generative model integrating single-cell time-frequency characteristics transformed from electrophysiological data with transcriptomic features

**DOI:** 10.1101/2024.03.29.587341

**Authors:** Kazuki Furumichi, Yasuhiro Kojima, Satoshi Nomura, Teppei Shimamura

**Affiliations:** Division of Systems Biology, Graduate School of Medicine, Nagoya University, Aichi, Japan; Laboratory of Computational Life Science, National Cancer Center Research Institute, Tokyo, Japan; Division of Computational and Systems Biology, Medical Research Institute, Tokyo Medical and Dental University, Tokyo, Japan

## Abstract

Patch-seq yields multi-modal data (e.g., transcriptomic and electrophysiological data) from single cells. However, currently used analytical methods are based on a few global electrophysiological features predefined from chronological potential changes and overlook the importance of time-frequency domain. In this study, we present LincSpectr, a deep neural network model that integrates transcriptomic and electrophysiological features with the latent variables of various variational autoencoders. This model combines the two modalities according to the connection between the latent variables of different modalities calculated by attention-like mechanisms and achieves cross-modal predictions and an inverse analysis. We discovered that the predicted electrophysiological features changed continuously along with their transcriptional profiles and that the neighborhood relationships between the latent states of the transcriptional profiles were consistent with those of the electrophysiological features. Inverse analysis of our model enabled the extraction of gene sets affecting specific time-frequency domains; some genes were likely to be involved in neural activity. Our approach represents a potential avenue to facilitate the discovery of molecular mechanisms underlying time-frequency activities in various cell types, enhancing our understanding of their roles in neural function.

## Introduction

The mammalian cerebral cortex is composed of neurons that exhibit diverse molecular, morphological, and electrophysiological features. These neurons have been classified into numerous cell types to understand complex brain circuits, and the recent advancements in single-cell transcriptomics have helped define them based on molecular markers^1^. In particular, Patch-seq, which links whole-cell patch clamping to high-quality single-cell transcriptomics, has been used as a powerful tool to clarify the comprehensive profiles of neuronal cell types^2^. The Patch-seq technique simultaneously measures gene expression and intracellular electrical activity and has identified several neuronal cell types (Pvalb, Sst, Vip, Lamp5, and three other excitatory classes)^3-5^. Previous studies have asserted that transcriptomic cell types are discrete and separable entities like chemical elements, but a recent study has shown that the morphoelectric distances between cell types is correlated with their transcriptomic distances^6^. This result suggests the possibility of a continuum of variation in morphoelectric types, which are related to transcriptomic variability.

To uncover the connections between disparate modalities such as transcriptomics (t-features) and intracellular electrophysiology (e-features), it is necessary to integrate single-cell multimodal data and quantify cross-modal correlations. Developments in multi-modal analysis techniques have enabled the extraction of gene sets linked to specific conductance and the prediction of cross-modal relevance based on Shapley values, among other achievements^7-9^. However, current methodologies have primarily focused on global electrophysiological characteristics, such as action potentials and spike frequencies, and cannot yield more localized information associated with transcriptomic effects, such as temporal components indicating neural activity decay and frequency components reflecting detailed activity pattern^10,11^.

Here, we introduce LincSpectr, a deep neural network model designed to integrate multimodal data by projecting t-features and e-features into their respective latent spaces using individual variational autoencoders (VAEs) and maximizing the similarity between latent variable pairs. This model learns from electrophysiological data transformed through continuous wavelet transform (CWT) and links the time-frequency characteristics with transcriptomic data; thus, enabling a nuanced analysis of the partial contributions of e-features. Additionally, by computing the Jacobian matrix for the average transcriptome of each cell type, we can infer the genes likely to be associated with these e-features. By capturing the continuous changes in e-features across cell types and linking them with transcriptomic variability, LincSpectr aims to elucidate the molecular pathways that define neuronal activity, potentially revolutionizing our understanding of cellular mechanisms in neuroscience.

## Results

### Conceptual view of LincSpectr

Fig. 1 describes the conceptual framework of LincSpectr. This method uses two heterogeneous types of data as inputs: t-features and e-features from single cells, as measured by Patch-seq. Here, “t-features” denote the count data matrix of gene expression in single neurons, and “e-features” refer to the time-frequency data matrix derived from each neuron’s membrane potential traces, transformed using CWT.

**Fig. 1:**
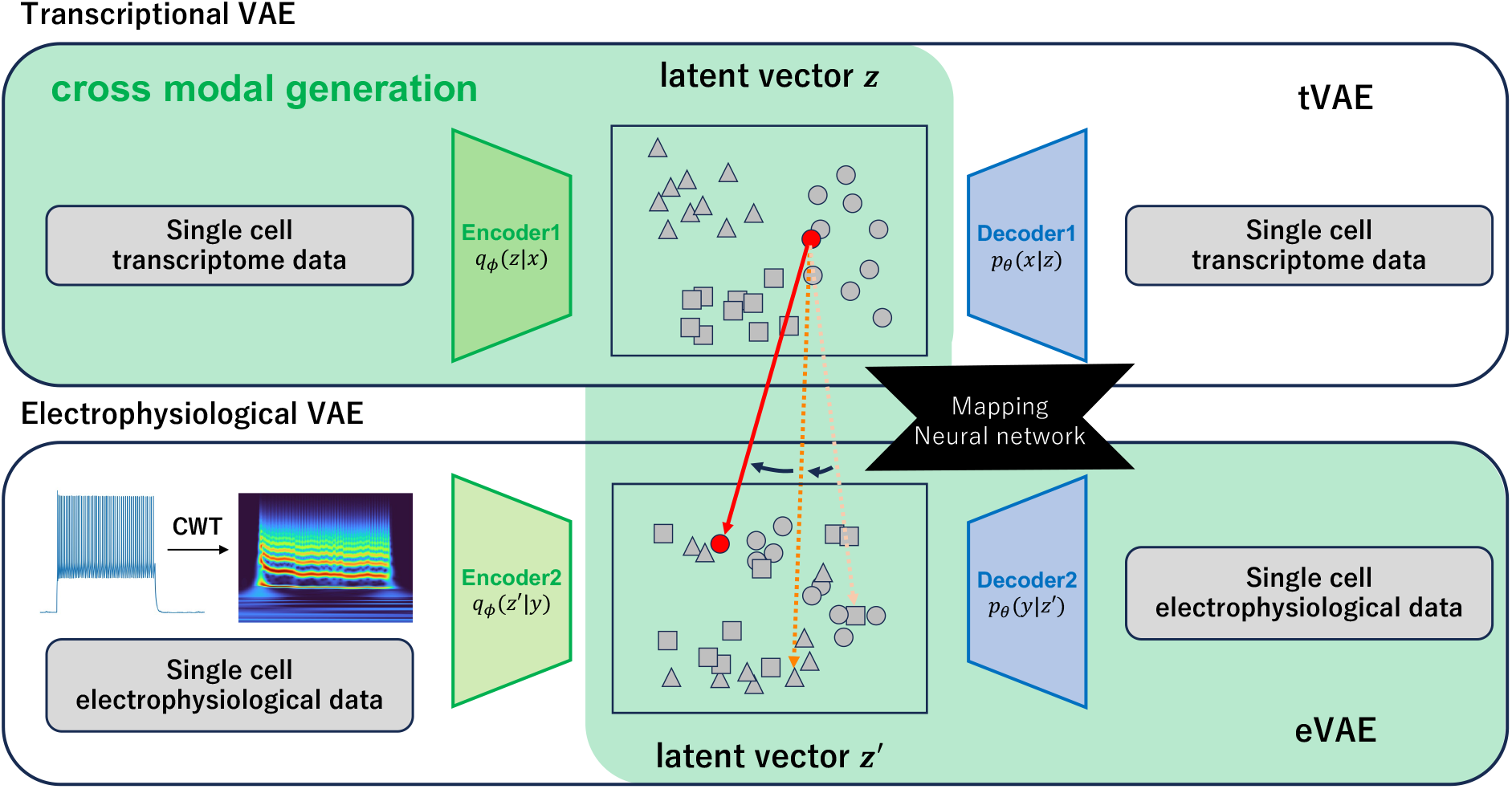
The architecture of LincSpectr integrating two modalities. Schematics showing the network architecture of LincSpectr based on the two latent spaces. They represent different modalities (transcriptomics and electrophysiology). The translation indicated with the red arrow conducted by the model maximizing the association between latent states of two modalities which were derived from an identical single cell. Leveraging this architecture, LincSpectr can take single-cell transcriptomic data as inputs and infer their electrophysiological data in the form of CWT.

LincSpectr uses two VAEs to achieve its objectives: an encoder maps t- and e-features to their respective independent latent spaces, whereas another decoder reconstructs the corresponding alternative modality from the variables within the latent space. The VAE for t-features is referred to as the tVAE and that for e-features is referred to as the eVAE. Additionally, LincSpectr integrates data across different modalities by maximizing the association between latent states of these modalities from the same cell through an attention-like mechanism. This integration elucidated the relationships between different data modalities, enabling a more comprehensive understanding of cellular functions.

Furthermore, an inverse analysis of the learned model, conducted using Jacobian analysis, revealed the effects of gene expression patterns on the e-features. This enabled the identification of continuous variations in e-features across cell types that are linked to changes in the transcriptome.

The advantages of our proposed method include its potential to significantly enhance our understanding of cellular molecular mechanisms and pave the way for new discoveries in neuroscience. LincSpectr is available as a user-friendly, open-source Python package at https://github.com/Kei0501/LincSpectr, offering a valuable resource for researchers seeking to explore and elucidate these complex cellular processes.

### Latent space representation of cell types and evaluation of cross-modal prediction accuracy

We used Patch-seq data from 1320 neurons derived from the adult mouse primary motor cortex and validated the clustering and cross-modal prediction accuracy of our proposed model. After excluding cells of insufficient quality (low read counts or excessive RNA contamination), 1228 cells were used for subsequent analysis. These cells were classified into nine transcriptomically defined cell types (t-types), including inhibitory classes (Pvalb, Sst, Vip, Lamp5, and Sncg) and excitatory classes (IT, ET, CT, and NP) using the Smart-seq2 and 10×sequencing technologies^6^.

By visualizing the latent spaces mapped by the tVAE and eVAE in the Uniform Manifold Approximation and Projection (UMAP), cells of the same type were appropriately aggregated in both modalities (Fig. 2a, 2b), except for CT neurons. Excitatory classes or neurons were not as clearly separated as the inhibitory neurons. This may be because glutamatergic cells form a largely continuous gradient of cells with correlated gradual changes in their expression profiles and cortical depths^12^.

**Fig. 2:**
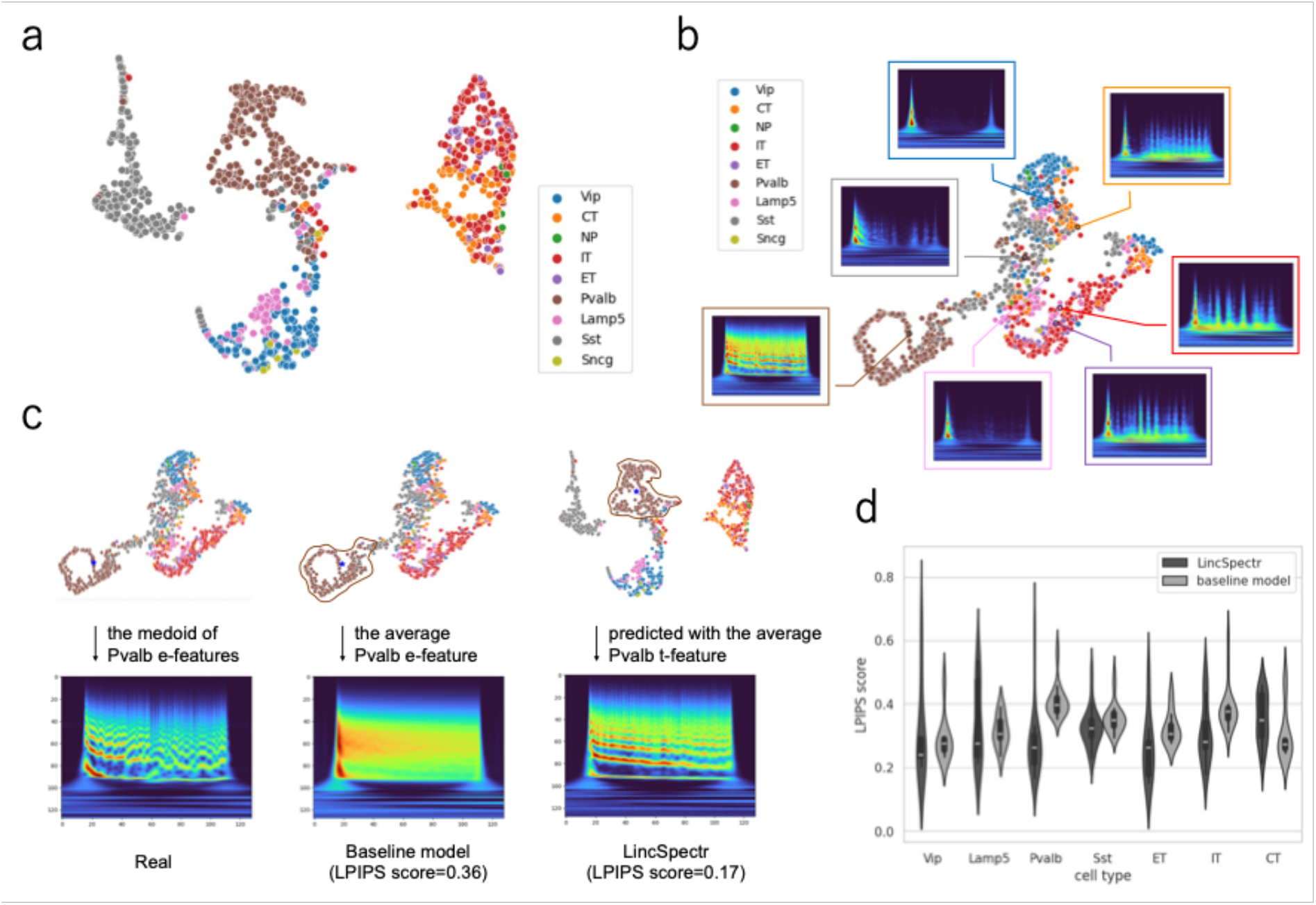
The latent representation and performance evaluation of LincSpectr. **a** The UMAP representation of the tVAE latent space of 1320 neurons from mouse motor cortex profiled with Patch-seq. The representation is colored by the transcriptomically defined cell types (t-types). The cells can be classified into nine cell types (five inhibitory classes: Pvalb, Sst, Vip, Lamp5, Sncg, and four excitatory classes: IT, ET, CT, NP). **b** The UMAP representation of the eVAE latent space colored by the t-types. The figure shows the medoid (indicated with a black circle) and average electrophysiological features (in boxes) of each population. **c** The comparison of the baseline model with LincSpectr for the representative Pvalb cell. The real e-feature of the medoid cell of the Pvalb cluster on the eVAE latent space (left), e-feature for the baseline model defined by the average e-feature of Pvalb cells and its Learned perceptual image patch similarity (LPIPS) score (middle), and e-feature predicted by LincSpectr from the average transcriptomic data of Pvalb cells (right). Each LPIPS score was based on the similarity between the real e-feature and the predicted e-feature. **d** The violin plot showing LPIPS scores for LincSpectr and the baseline model in the task of predicting e-feature for each cell type (**c**). This represents a distribution two ways: a patch showing a symmetric kernel density estimate, and the quartiles/whiskers of a box plot.

Next, we reconstructed the e-features of the medoid cells (cells with the minimum sum of distances to all other cells in the cluster) in the latent space of the eVAE for the seven major cell types. The reconstructed time-frequency patterns showed that signals increased across a broad range of frequencies at the beginning of activity, and weak signals were observed at the end for all cell types (Fig. 2b). Pvalb cells exhibit characteristic activation patterns across several frequency domains over long periods. E-features of ET, IT, and CT cells, classified as excitatory neurons, showed relatively similar characteristics in that signals alternated between enhancement and attenuation during activity.

To examine the cross-modal prediction accuracy of LincSpectr, we predicted e-features using transcriptome data from 132 cells that were excluded from training in advance. We created a baseline model for comparison, which was the pixel average of all e-features of the cells classified into each t-type. The Learned Perceptual Image Patch Similarity (LPIPS) score^13^ was used to quantitatively evaluate the predictive performance. The LPIPS score can be used to calculate the similarity between two images using a predefined network, and its accordance with human perception has already been demonstrated. The LPIPS scores for average t-features of Pvalb cells predicted by each model were 0.17 for LincSpectr and 0.36 for a baseline model; hence, demonstrating the high prediction accuracy of LincSpectr (Fig. 2c). The median LPIPS score of LincSpectr exceeded that of the baseline model in six out of seven cell types (Fig. 2d). Among the seven cell types, Pvalb, Sst, and IT neurons showed statistically significant differences in LPIPS scores.

These results show that LincSpectr is highly effective in conducting precise analysis and generating cross-modal predictions based on cell types.

### Clustering based on e-features and their relationship with gene expression profiles

To investigate the type of e-features that separated the cells in the latent space of the eVAE, cells were clustered into seven clusters, which we denote as e-types in this article, using the *k*-means algorithm without prior information on cell types. Furthermore, the e-features of the cells in each cluster were averaged, and the e-features characterizing each cluster were visualized (Fig. 3a, b). We discovered that the pairs of Pvalb and Sst cells, Vip and some Sst cells, IT and ET cells, as well as Lamp5 and some Sst cells were grouped into identical clusters, indicating that these pairs exhibited consistent e-features.

**Fig. 3:**
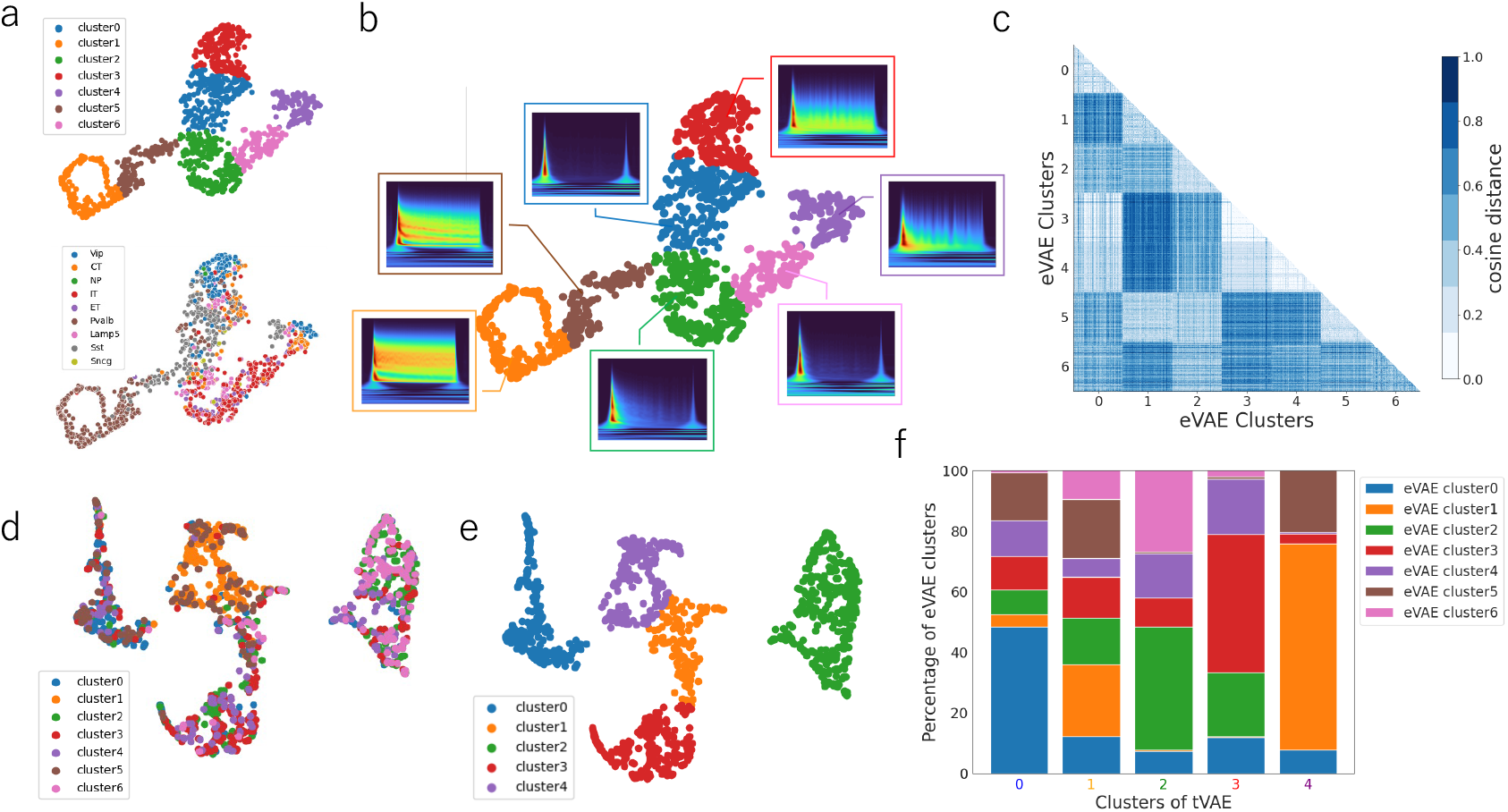
Classification of e-types and their relationship to t-types. **a** The UMAP representation of the eVAE latent space, colored by the cluster labels determined by the *k-means* algorithm (upper). The same UMAP representation colored by t-types (lower). **b** The average electrophysiological features (in boxes) of each cluster defined in (**a**). **c** The heatmap showing the values of cosine distances between the cells. This figure only represents the results for the 700 cells, 100 cells extracted from each of the 7 clusters. **d T**he UMAP representation of tVAE latent space colored by the cluster labels as in (**a**). **e** The clusters on the tVAE latent space determined by the *k-means* algorithm. The cells are classified into five clusters and colored by cluster labels. **f** 100 percent stacked bar graph showing the ratio of e-types for each tVAE cluster.

In addition, we extracted 100 cells from each cluster and represented their cosine distances on a heat map. The values of cosine distance were small not only between the cells from the same cluster, but also between the cells from adjacent clusters (e.g., clusters 0 and 3, 1 and 5, 2 and 5, and 2 and 6) (Fig. 3c). Thus, adjacent clusters on UMAP are likely to show similar e-features and are spread over the same cell-type population. This suggests that e-features are not independent of each type but rather vary continuously with gene expression.

To determine the relationship between e-types and t-types, we examined cell-type annotations based on the gene expression profiles of the cells in each type. Cluster 1 contained ∼96.5% Pvalb cells (191/198 cells), and cluster 6 contained ∼84.6% IT cells (104/123 cells). These clusters were dominated by a single cell type. In contrast, clusters that included two cell types were observed equally. Cluster 4 contained ∼29.2% CT cells (40/137 cells) and ∼23.4% Vip cells (32/137 cells), whereas cluster 5 contained ∼59.0% Pvalb cells (69/117 cells) and ∼39.3% Sst cells (46/117 cells). The other clusters had one dominant cell type. Cluster 0 had ∼65.4% Sst cells and CT cells >20%, cluster2 had ∼48.6% IT cells and ET, Lamp5, and Sst cells >20%, whereas cluster 3 had ∼53.2% Vip cells and CT and Sst cells over >20%. This suggests that some groups of cells exhibited consistent e-features among the cell types. The heterogeneity of the e-features can be associated with the finer subtypes of each cell type^8^. Moreover, we discovered that both excitatory and inhibitory cell types were included in the same cluster, consistent with the findings from a previous study^11^; for these cell types, we observed continuous changes in the e-features of glutamatergic and GABAergic neurons in the primary motor cortex of mice.

Furthermore, we displayed the classified cluster information of the eVAE onto cells in the tVAE latent space and observed that each type was aggregated in the tVAE latent space. The cells in the tVAE latent space are classified into five clusters, denoted as t-types, using *k*-means algorithm. Next, we analyzed the overlap between e-types and t-types. The cells in cluster 0 were derived from various clusters in the eVAE latent space, whereas the other clusters contained nearly half of their cells from a single cluster. Approximately 40.7%, 48.4%, 68.2%, and 45.8% of the cells belonged to specific clusters in the eVAE for clusters 1, 2, 3, and 4, respectively. Reflecting on the type labels of the cells in the tVAE latent space, we confirmed that numerous cells in the cluster were derived from clusters with close e-features. This indicates that the classification based on e-features is also valid in the tVAE latent space (Fig. 3d–f). In contrast, exceptional clusters with distinctly different e-features correspond to regions in the tVAE latent space where different cell types are mixed. Because Patch-seq recordings often contain contaminating mRNA from neighboring neurons or non-neuronal cells,^11^ the quality of these cells may have been insufficient for analysis.

### Genes affecting e-features of each cell type

The standard e-features of each cell type may be influenced by the differences in the expression patterns of specific genes. For instance, cortical neurons expressing Pvalb genes show the e-feature of a high-frequency firing rate, which is determined by membrane potential changes associated with their types, expression levels, and ion channel densities^14^. Thus, to elucidate the molecular mechanisms and biological pathways underlying e-features, an inverse analysis of the model was performed using a Jacobian matrix. We extracted genes with large contributions to e-feature prediction by singular value decomposition of the Jacobian matrix and conducted gene set enrichment analysis using Enrichr. In addition, we visualized the time-frequency features corresponding to the extracted singular values to show how the fluctuation of each gene was associated with the e-features.

The analysis showed that, for all cell types, genes with large contributions were associated with Gene Ontology (GO) terms related to neuronal activity. For example, GO terms strongly associated with contributing genes in Vip cells include sodium: phosphate symporter activity such as cellular glutamate transporters. The contributing genes in Pvalb cells were enriched in GO terms related to calcium ion binding, indicating a relationship with a calcium binding protein (CABP5), which depolarizes membrane potential by inhibiting the inactivation of L-type calcium channels ^15^. The contributing genes in ET and IT cells showed significant overlap with GO terms related to ligand-gated channels, such as acetylcholine receptors, which are likely to affect various channels and transporters, including inward-rectifying potassium channels. GO terms enriched in Lamp5 cells are associated with PDZ domain binding, whereas those in Sst cells are related to secondary active transmembrane transporter activity.

Several genes that markedly contribute to cross-modal prediction have been reported to be involved in neuronal function or differentiation. Tbata, the levels of which exhibited large changes in inhibitory classes of neurons (Fig. 4a–c), is predicted to interact with Kinesin Kif17, affecting dendritic transport, and is implicated in neurite outgrowth and dendritic patterning via the nerve growth factor signaling pathway^16^. Gpr6, which showed high contribution in Pvalb cells (Fig. 4c), is a type of G protein-coupled receptor that has been reported to promote neurite outgrowth by positively regulating intracellular cAMP in neurons^17^. Chrna5, which plays an important role in both ET and IT cells (Fig. 4e, f), promotes the expression of nicotinic acetylcholine receptors and facilitates the introduction of postsynaptic cholinergic responses^18^. While genes whose neuronal functions have been previously identified were detected, genes linked to the immune system or those with unclear functions were also extracted. This suggests that genes that have not received much attention might affect the e-features.

**Fig. 4:**
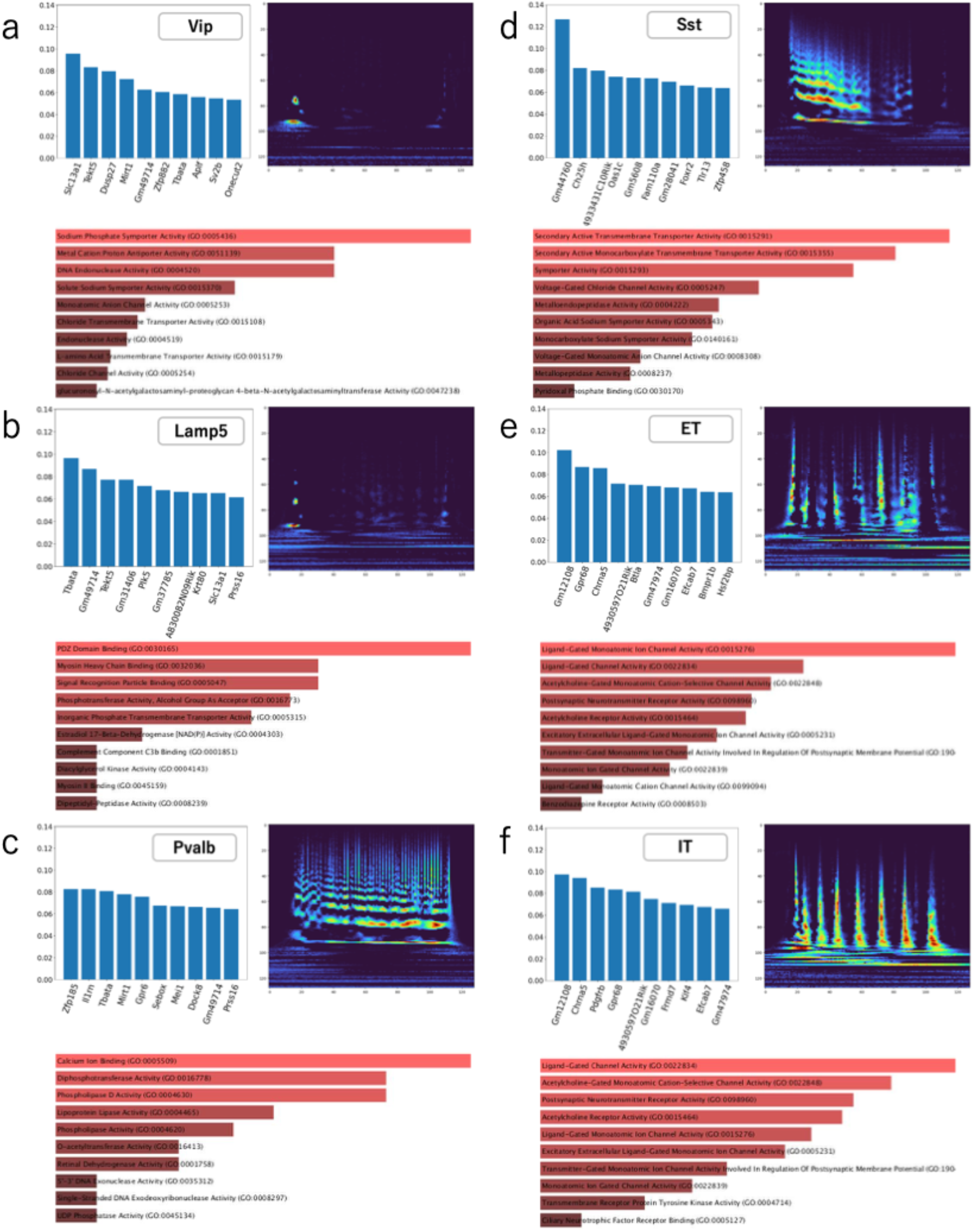
Identification of gene sets contributing to e-features of each cell type. **a-f** Results of the inverse analysis of LincSpectr for each cell type. Each figure indicates the result for Vip cells (**a**), Lamp5 cells (**b**), Pvalb cells (***c***), Sst cells (***d***), ET cells (**e**), and IT cells (**f**). The figure shows the genes with large values in singular value decomposition of the Jacobian matrix (upper left), the time-frequency features corresponding to the singular values (upper right), and GO terms enriched in each cell type (lower). In the lower figures, the length of the bars indicates the importance of that particular GO terms (based on their p-values).

## Discussion

In the present study, we used two VAEs to develop LincSpectr, a model that enables cross-modal analysis between single-cell transcriptomics and electrophysiological data by leveraging probabilistic distributions. Furthermore, the inverse analysis using vector Jacobian products allowed us to predict the genes that contributed to variations in electrophysiological features. LincSpectr uniquely elucidates the relationship between the time-frequency aspects of electrophysiological characteristics and transcriptomics through the application of CWT. This approach is particularly valuable for investigating neurons that demonstrate “resonance,” a phenomenon in which certain frequencies of electrical input are preferentially amplified^19^. Such time-frequency-focused models hold promise for uncovering the molecular mechanisms underlying frequency-dependent responses. Our analysis validated that the latent spaces for each modality accurately represented the cellular features, demonstrating superior predictive performance over baseline models across most cell types. Additionally, by analyzing the Jacobian matrix, we were able to identify genes likely to impact specific time-frequency electrophysiological characteristics.

Furthermore, Jacobian analysis enabled us to explore the time-frequency domains that were predicted to be altered by the perturbation of the expression of each gene. The high signals in the Jacobian analysis observed at the beginning and end of the time components of the e-features of all cell types corresponded to the onset and offset timing of the stimuli, respectively. Patterns observed in the analysis of Pvalb cells showed high-frequency spike firing encompassing multiple frequencies with high amplitudes during neuronal activity, suggesting an association with the short-term facilitation of synaptic transmission. Calcium ions can affect this activity, with Gpr6 potentially activating the cAMP/PKA signaling pathways to induce rapid firing by controlling the downstream expression of voltage-dependent Ca^2+^ and voltage-dependent Na^+^ channels^17^. Early activity patterns observed in Vip and Lamp5 cells indicate initial bursts followed by activity suppression after stimuli, implying that they possess characteristics of early adaptation. The top genes included Tbata, which is known to be involved in neuronal morphogenesis during dendrite and axon formation^16^. The initial bursts may have been influenced by differences in cellular morphology. ET and IT cells showed sustained and slightly delayed spike-firing patterns, which were likely to be associated with ligand-gated channels. Ligand-dependent ion channels, such as Chrna5, control membrane capacitance and input resistance, including the expansion of the action potential width unique to excitatory neurons^18^.

This study introduced a model that connects transcriptomic and electrophysiological features from Patch-seq data, a methodology potentially extendable to other modalities. The inclusion of morphological data in Patch-seq enables the prospective construction of a VAE for morphology, predicting the impacts of various modalities on morphological data. Our approach focuses on e-features during the intermediate phases of multiple stimuli, potentially overlooking long-term characteristics. We will address this limitation through a comprehensive analysis of larger datasets. Notably, the absence of predicted potassium channels, such as HCN1^20^, in our inverse analysis was unexpected. This discrepancy may stem from the challenge associated with distinguishing the influence of genes expressed across all cell types. To mitigate partial overfitting and enhance the generalizability of the model, further pre-training with an expanded Patch-seq dataset^21^ is necessary; however, acquiring substantial Patch-seq data remains a challenge. This problem can be solved by using patch clamp data and sc-RNAseq data. Besides, overfitting risks could be avoided by adding dropout layers or introducing noise into the data at a certain probability like denoising auto encoder^22^.

As technologies for obtaining multi-modal data from single cells have advanced, the demand for multi-modal analysis has significantly increased. Our study proposes a systematic framework for elucidating the correspondence between transcriptomic and electrophysiological features, offering an efficient tool for understanding the molecular underpinnings of the diverse neural networks in the brain and identifying new molecular targets for experimental exploration.

## Methods

### Model structure

We used VAEs^23^ for two modalities (t-features and e-features) and combined them with a model to learn the probabilistic relationship between the two types of latent variables and achieve cross-modal predictions. The model consists of a tVAE encoder, a model to predict the correspondence between the tVAE and eVAE, and an eVAE decoder. We trained the two VAEs, and then optimized the parameters of the connection model.

In the VAEs, we assumed a prior distribution *p*(*z*) for the latent variable *z* and a conditional probability distribution *p*_θ_(*x*|*z*), which have optimizable parameter *θ* for an observed variable *x*. Then, we approximated the posterior distribution *p*_*θ*_ (*z*|*x*), which is difficult to calculate, using a variational posterior distribution *q*_*φ*_(*z*|*x*), which have optimizable parameters *φ* by variational inference. To optimize the parameters of the model that allow it to produce estimates, we maximize the evidence lower bound (ELBO), which lead to maximizing *log* (*p*_*θ*_ (*x*)) and minimizing *D*_*KL*_[*q*_*φ*_ (*z*|*x*)||*p*_*θ*_(*z*|*x*)] simply and simultaneously, which are difficult to calculate separately. ELBO was defined as:

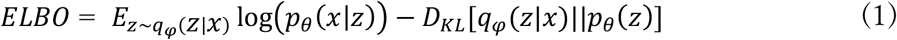

The first term on the right side is the reconstruction error shows the agreement between the input and generated data. The second term is the Kullback–Leibler divergence (KL divergence) expression indicates the distance between the approximated posterior and prior distributions. Here, the prior distribution *p*(*z*) are assumed to be a standard multivariate normal distribution and the approximate posterior distribution *q*_*φ*_(*z*|*x*) are assumed to be Gaussian distributions *q*_*φ*_ (*z*|*x*) = *N*(*z*|*μ*_*φ*_ (*x*),*diag*(*σ*_*φ*_^2^(*x*)) with parameters *μ*_*φ*_ (*x*) and *diag*(*σ*_*φ*_^2^(*x*)) generated by the encoder neural network. This sampling was computed *L* times to minimize the loss function expressed in Equation (2), whose expected value coincided with the negative ELBO by stochastic gradient descent. For the reconstruction error, we conducted a Monte Carlo approximation by sampling *z* on *L* times from the variational posterior distribution.

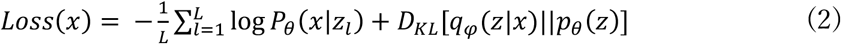

Because the first term in equation (2) has no gradient with respect to the parameter *φ* of the variational distribution *q*, a reparameterization method was used to allow back propagation. The reparameterization method implements sampling by computing 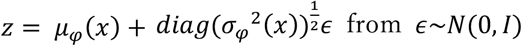 from *ϵ∼N*(0, *I*), making the objective function stochastic and providing a solution for the gradient. This allows the optimization of Equation (2) through backpropagation.

### (1) tVAE structure

The first model using this VAE received data of t-types *X*_*t*_ *∈ ℕ*^*C×G*^ (*C* cells and *G* genes) as input and decoded data of same array dimension from the encoded 10-dimensional latent variable *z*^(*t*)^ . The encoder consists of two LinearReLU layers, which combine linear transformations and ReLU activation, and two linear coupling layers. The decoder consists of two LinearReLU layers and one linearly coupled layer, with the output layer being activated by Softplus. *X*_*t*_ is the gene count data; therefore, a generation model using the Poisson distribution was employed as the first term of the ELBO. The conditional probability of the generation based on the Poisson distribution was expressed as:

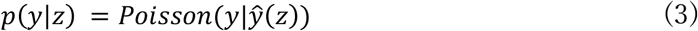

Minimizing this loss function, we optimized the *qφ* (*z*^(*t*)^|*X*_*t*_) and *p*_*θ*_ (*X*_*t*_|*z*^(*t*)^) equations.

A small value (1.0 ×10^-10^) was added to *ŷ* at each epoch because some elements of *ŷ* (*z*) showed extremely small values during training.

### (2) eVAE structure

The second model using VAE received e-types data *X*_*e*_ *∈ ℕ*^*C×E*^ (*C* cells and *E* electrophysiological features) as input and was constructed as the first VAE model. The electrophysiological features *E* were transformed by CWT as described below (subheading “Dataset preprocessing”). CWT is a method waveform data analysis in which the frequency spectral change of the Fourier transform is represented in two dimensions simultaneously with a temporal or spatial transition. This method enables the spectral analysis of sudden and nonstationary waveform temporal changes. The difference from the tVAE is that we added one additional layer of LinearReLU to the decoder and activated the output layer using a sigmoid. In addition, we used *p*(*X*_*t*_*Vz*^(*t*)^) = *𝒩*(*X*_*t*_|*μ*^(*t*)^(*z*^(*t*)^, *σ*^2^*I*)) as the loss function. The optimization of *q*_*φ*_ (*z*^(*e*)^|*X*_*t*_) and *p*_*θ*_ (*X*_*t*_|*z*^(*e*)^) was achieved by minimizing the loss function.

### (3) Connection model structure

To link these two VAEs, a model was developed to estimate the corresponding latent variable *z*^(*e*)^ of the eVAE from the latent variable *z*^(*t*)^ of the tVAE. The strength of the association between the two latent variables was assumed to be expressed as:

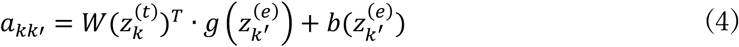

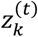 is the latent variable of tVAE for cell *k* and 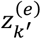 is the latent variable of eVAE for cell *k*^*′*^. The probability of observing 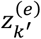 from 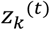 can be written as the following formula for calculating the *P*_*kk′*_:

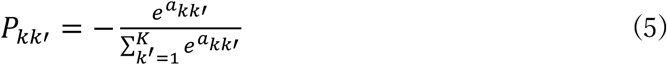

Here, *K* denotes the total number of cells to be considered. This was computed by applying Softmax to *a*_*kk′*_. By maximizing *P*_*kk*_ and optimizing the parameters of *W, g*, and *b*, the model can output 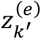, which allowed us to predict 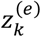 from the corresponding 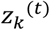.

We created a list of *z*^(*e*)^ values generated by the second VAE and prepared an inference model that outputs the Softmax probability distribution for each cell state by maximizing Equation (4) for the *z*^(*t*)^ input to the model.

By combining this inference model with a tVAE encoder and an eVAE decoder, a model that outputs *X*_*e*_ in response to *X*_*t*_ input was generated.

### Training procedure

All the optimizers used Adam to train the model. Each model was trained on 85% of the data, with 10% used for validation and 5% used as test data. The tVAE was trained with minibatch sizes of 16 and 350 epochs and a learning rate of 0.004, whereas the eVAE was trained with minibatch sizes of 16 and 300 epochs and a learning rate of 0.0001.

The model used to estimate the latent space was trained with mini-batch sizes of 512 and 350 epochs and a learning rate of 0.0001, splitting the dataset in similar proportions.

### The inverse analysis of the model

To predict the elements of the t-types that deform the e-types, we performed an inverse analysis of the learned model using a Jacobian matrix of e-features differentiated by t-features. With the sequence length of t-type *G* and the sequence length of e-type *E*, the Jacobian matrix was expressed as:

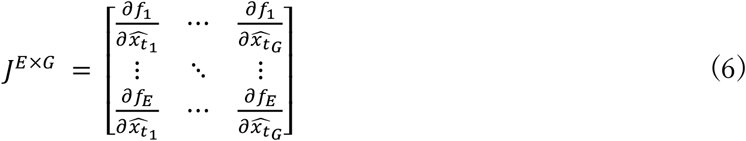

Furthermore, singular value decomposition was performed on the Jacobian matrix to extract the principal perturbation patterns of e-features by t-features.

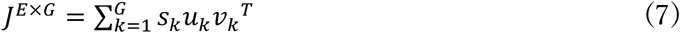

where *G* < *E, u*_*k*_ and *v*_*k*_ are the eigenvectors of the target matrix, indicating the data corresponding to the *k*^th^ largest singular value. Thus, by analyzing a higher *v*_*k*_, we can extract the molecules that affect the e-features corresponding to *u*_*k*_.

## Benchmarking and evaluation

### Dataset preprocessing

For our analysis, we downloaded the Patch-seq dataset of adult mouse motor cortex published by Scala et al^6^. The t-type data were obtained from https://github.com/berenslab/mini-atlas, and the data for e-types were obtained from the DANDI archive, https://dandiarchive.org/dandiset/000008.

Next, the t-types, represented by read count data for exons, were extracted from single neuron using Smart-seq2 and recorded in m1_patchseq_exon_counts.csv. In a single cell, 42466 genes including pseudogenes and annotated non-coding regions, were detected.

To use these data for analysis, genes with high variability in expression were selected and gene expression was quantified. We used Scanpy, a toolkit for analyzing single-cell gene expression data, and removed cells expressing fewer than 100 genes and genes expressed in fewer than 10 cells. To compare the values among cells, we normalized each cell by total counts over all genes (target_sum=1e4) and log-transformed the count data. Subsequently, genes were filtered by identifying the top 2000 genes based on mean expression levels and variance using the highly_variable_genes function.

E-types are the membrane potential traces per cell, as recorded by Patch-seq. The data were recorded from neurons injected with 600-ms-long current pulses; the current was increased from -200 pA to 1380 pA in 20-pA increments. We extracted the 30^th^ data point from the beginning of the membrane potential trace and performed CWT using ssqueezepy (https://github.com/OverLordGoldDragon/ssqueezepy). For the processed data, a scale of time components corresponded to 1/25000 seconds, and that of frequency indicated specific frequencies calculated by ssqueezepy for an e-feature previously decided.

For signals in the high-frequency domain that could be noise, we checked the data for 4 ms from the beginning of the recording. We removed the signals in the high-frequency regions (over 200 among the 277 classified frequencies) that did not fall within the average ±1 SD and compressed the data in the time direction. To reduce the computational volume, the resolution was reduced from 277×25000 to 128×128 after removing the regions where the signals were hardly observed (75/277 of the low-frequency component and 1/5 of the time component).

The t-types of 1329 cells were practicable, and 1320 cells were used for training, excluding cells with no information on paired e-types or cells with fewer than 30 membrane potential traces.

### UMAP visualization of the transcriptomic and electrophysiological data

To visually understand how information is represented in the latent space of the VAE, a dimensionality reduction of the data was performed using UMAP. UMAP is grounded in algorithms based on topological data analysis, which captures datasets of continuous structures in high-dimensional space and projects them onto low-dimensional spaces^24^. For each VAE, 10-dimensional latent variables were transformed with UMAP and displayed in a scatter plot in a 2D space. We set the parameter n_neighbors (limiting the size of the local neighborhood) to 30 and min_dist (indicating the minimum distance allowed for a point in the low-dimensional representation) to 0.01.

To color-code each point by cell type, the exon count data m1_patchseq_exon_counts.csv and patch-seq metadata m1_patchseq_meta_data.csv were combined using cell names providing cell type information.

### LPIPS score calculation

The LPIPS score was used to evaluate the performance of the model^13^. The LPIPS score represents the distance between images and is calculated based on features acquired by the convolution layer of a learned image classification network, such as AlexNet or VGG, for the input images. Compared to methods that focus on pixel luminance and contrast, the LPIPS score quantifies errors with a precision closer to that of human perception. The errors in pixel shifting can be properly evaluated. After loading the trained network, the calculation was performed by inputting the two torch-type images for comparison.

### Performance comparison with the baseline model

We compared the performance of the baseline model with that of the reconstructed data by calculating the LPIPS scores for the correct data. For the baseline model, the dataset used as validation data was classified by cell types, and the average images of the e-features were used as the output. The reconstructed data from LincSpectr were used to predict the e-features of medoid cells. The correct data are the e-features for the medoid cells of the cluster of target cell types in the eVAE.

### Comparison of similarities among clusters

To determine the similarities of e-features from each cluster in the eVAE latent space, the cosine distance (a measure of similarity between two nonzero vectors defined in the inner product space) was calculated and displayed on a heatmap. The cells of clusters in the eVAE latent space classified using the *k*-means algorithm were compared.

### Identification of gene sets associated with electrophysiological features

The inverse analysis was performed on the model after training (subheading “The inverse analysis of the model”). The higher the singular value, the greater the impact on the Jacobian matrix; hence, we analyzed the components with large singular values. We extracted *v*_*k*_(*k* = 0 or 1) corresponding to the first or second singular values and obtained 2000-dimensional gene variation data. The genes included in the top 10 are shown in the bar graph as those that strongly affect the cell type (Fig.4a-f). The top 50 genes in *v*_*k*_(k = 0 or 1) were also extracted and GO analysis was performed using Enrichr. We used the gene-set library “GO Molecular Function 2023” for this analysis. The length of the bars indicated the importance of that particular GO terms (based on their p-values). The same is true for the brightness of the color; the brighter the color, the greater the importance of that term. To confirm that the time-frequency component was affected by variations in these genes, *u*_*k*_(*k* = 0 or 1) was displayed in a heatmap as 128×128 images.

## Acknowledgements

This work was supported by a grant from Moonshot R&D (T.S., grant number JPMJMS2025) from the Japan Science and Technology Agency (JST).

## Author contributions

Y.K. conceived the concept of the method. K.F. designed the source code for this method, conducted experiments to verify its validity, designed the analysis using this method, and performed the analysis under the supervision of Y.K. and T.S. S.N. made minor modifications to the theory of this method. All the authors have read and approved the final version of the manuscript.

## Competing interests

The authors declare no competing interests.

## Code and Data Availability

The LincSpectr implementation is available at https://github.com/Kei0501/LincSpectr. The t-type dataset were obtained from https://github.com/berenslab/mini-atlas, and the dataset for e-types were obtained from the DANDI archive, https://dandiarchive.org/dandiset/000008.

## Notes

https://github.com/Kei0501/LincSpectr

https://github.com/berenslab/mini-atlas

https://dandiarchive.org/dandiset/000008

### Competing Interest Statement

The authors have declared no competing interest.

